# Deep reinforcement learning for the control of microbial co-cultures in bioreactors

**DOI:** 10.1101/457366

**Authors:** Neythen J. Treloar, Alexander J.H. Fedorec, Brian P. Ingalls, Chris P. Barnes

**Affiliations:** Department of Cell and Developmental Biology, University College London, WC1E 6BT, UK; Department of Applied Mathematics, University of Waterloo, N2L 3G1, Canada; UCL Genetics Institute, University College London, WC1E 6BT, UK

**Keywords:** reinforcement learning, chemostat, bioreactor control, artificial intelligence

## Abstract

Multi-species microbial communities are widespread in natural ecosystems. When employed for biomanufacturing, engineered synthetic communities have shown increased productivity (in comparison with pure cultures) and allow for the reduction of metabolic load by compartmentalising bioprocesses between multiple sub-populations. Despite these benefits, co-cultures are rarely used in practice because control over the constituent species of an assembled community has proven challenging. Here we demonstrate, *in silico*, the efficacy of an approach from artificial intelligence – reinforcement learning – in the control of co-cultures within continuous bioreactors. We confirm that feedback via reinforcement learning can be used to maintain populations at target levels, and that model-free performance with bang-bang control can outperform traditional proportional integral controller with continuous control, when faced with infrequent sampling. Further, we demonstrate that a satisfactory control policy can be learned in one twenty-four hour experiment, by running five bioreactors in parallel. Finally, we show that reinforcement learning can directly optimise the output of a co-culture bioprocess. Overall, reinforcement learning is a promising technique for the control of microbial communities.

## 1 Introduction

The ability to engineer cells at the genetic level has enabled the research community to make use of biological organisms for many functions, including the production of biofuels [1, 2, 3], pharmaceuticals [4] and the processing of waste products [5]. Communities consisting of multiple strains of cells have been shown, in some cases, to be more productive than monocultures at performing processes such as biofuel production [2, 3, 6] and alleviate the problem of metabolic burden that occurs when a large pathway is built within a single cell [7]. For these reasons co-cultures should play a significant role in the advancement of bioprocessing. However, maintaining a co-culture presents its own set of problems. The competitive exclusion principle states that when multiple populations compete for a single limiting resource, a single population with the highest fitness will drive the others to extinction [8]. It has been proven that, under ideal conditions, at most one population can survive indefinitely in a chemostat where multiple cell populations are competing for a single substrate [8]. An additional challenge is that the interactions between different populations of microbes can make long term behaviour in a co-culture difficult to predict [9]; the higher the number of distinct populations, the greater the challenge becomes to ensure system stability [10].

Previously, methods of co-culture population control have been engineered into cells genetically, e.g. using predator-prey systems [11] or mutualism [7, 12]. However, processes such as horizontal gene transfer and mutation make the long term genetic stability of a population hard to guarantee [9], meaning that genetic control methods can become less effective over time. Another potential problem is the increased metabolic load imposed on a cell due to the control genes, which can leave less resources for growth and the production of useful products [13]. These downsides can be avoided by exerting control over the environment, which is the dominant approach in industry. Established techniques are Proportional-Integral-Derivative controllers [14], Model-Predictive-Controllers [15, 16, 17] or the development of *ad hoc* feedback laws [18, 19, 20, 21]. Here we investigate the viability of reinforcement learning as a complement to these methods.

For our analysis, we use the chemostat model, which provides a standard description of bioprocess conditions. This model is applicable to wide range of other systems where cell or microorganism growth is important, including wastewater treatment [22] and the gut microbiome [23]. Such systems can be especially difficult to control because they often are equipped with minimal online sensors [24], limiting the effectiveness of classical control techniques that are hampered by infrequent or delayed system measurements [20, 25].

Reinforcement learning is a branch of machine learning concerned with optimising an agent’s behaviour within an environment. The agent learns an optimal behaviour policy by observing environmental states and selecting from a set of actions that change the environment’s state. The agent learns to maximise an external reward that is dependent on the state of the environment. The training of a reinforcement learning agent is often broken up into *episodes*. An episode is defined as a temporal sequence of states and corresponding actions (generated by the agent interacting with the environment) until a terminal state is reached. The total reward obtained during an episode is called the return. For this study, we used a data-efficient variant of reinforcement learning called Neural Fitted Q-learning [26, 27, 28] (see Methods).

Much reinforcement learning research has been done on video games [29] due to the availability of plentiful training data. However, it is also seeing application to more practical problems in the sciences, including to optimise chemical reactions [30] and in deriving optimal treatment stategies for HIV [31] and sepsis [32]. A partially supervised reinforcement learning algorithm has also been applied to a model of a fed-batch bioreactor containing a yeast monoculture [33].

Here we develop a control scenario in which the growth of two microbial species in a chemostat is regulated through the addition of nutrients *C*_1_ and *C*_2_ for which each species is independently auxotrophic (Figure 1A, 1B). The influx of each nutrient is controlled in a simple, on-off manner (bang-bang control). At each time point, the agent decides, for each auxotrophic nutrient, whether to supply this nutrient to the environment at the fixed inflow rate over the subsequent inter-sample interval. This constitutes the set of possible actions. A constant amount of carbon source, *C*_0_, is supplied to the co-culture. We define the system state as the population levels of each population in the chemostat (assumed to be measured using fluorescence techniques). The objective is either to maintain specific population levels or to maximize product output. A corresponding reward is given, which depends on the distance of the population levels from the target value or as a function of product output. The populations evolve continuously, and the reward is likewise a continuous function of the state. In contrast, the agent’s actions are discrete (bang-bang), and are implemented in a sample-and-hold strategy over a set of discrete sampling times. A visual representation of a two-population case is shown in Figure 1C.

**Figure 1:**
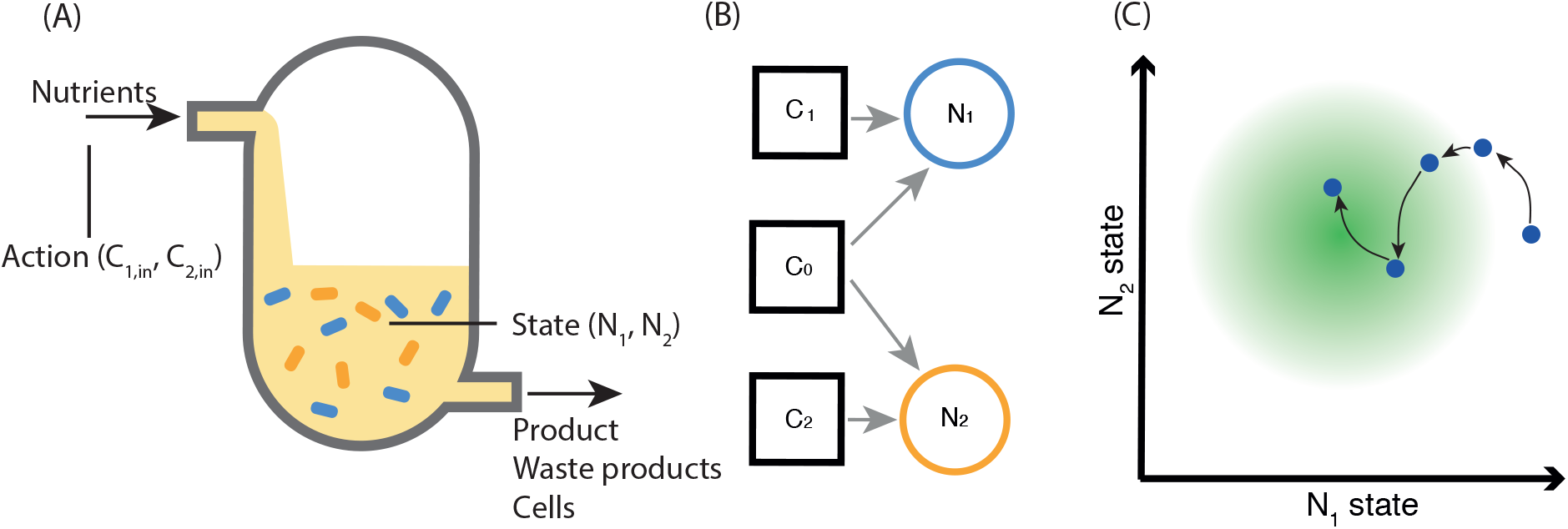
(A) Diagram of a chemostat. The state observed by the reinforcement learning agent is composed of the populations of two strains of bacteria; the actions taken by the agent control the concentration of auxotrophic nutrients flowing into the reactor. (B) System of two auxotrophs dependent on two different nutrients, with competition over a common carbon source. (C) Representative system trajectory. The agent’s actions, taken at discrete time-points (circles), influence the state dynamics (red arrows), with the aim of fulfilling the reward condition (moving to the centre of the green circle). The state is comprised of the (continuously-defined) abundance of two microbial populations, **N**_1_ and **N**_2_. The agent’s actions dictate the rate at which auxotrophic nutrients flowing into the reactor. At each time-step, the agent’s reward is dependent on the distance between the current state from the target value.

Below, we illustrate that an agent can successfully learn to control the bioreactor system in the customary episodic manner. Secondly, we compare a reinforcement learning approach to proportional integral control, both working in a model free way on simulated data, and show that the learning approach performs better in situations where sampling is infrequent. We then show that the agent can learn a good policy in a feasible twenty four hour experiment. Finally, we demonstrate that reinforcement learning can be used to optimise productivity from direct observations of the microbial community. Traditional proportional integral control could only be applied to such a case via a model of the system, or with additional measurement data from further online sensors.

## 2 Results

### 2.1 A mathematical model of interacting bacterial populations in a chemostat

We develop a general model of *m* auxotrophs growing and competing in a chemostat. The model captures the dynamics of the abundance of each species (*m*-vector **N**), the concentration of each auxotrophic nutrient (*m*-vector **C**), and the concentration of the shared carbon source (scalar *C*_0_). A sketch of the two-species case is shown in Figure 1 A-B.

The rate of change of the concentration of the shared carbon source is given by:

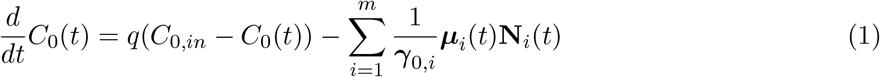

where *γ*_0_ is a vector of the bacterial yield coefficients for each species, *C*_*0,in*_ is the concentration of the carbon source flowing into the bioreactor, *μ* is the vector of the growth rates for each species, and *q* is the flow rate.

The concentration of each auxotrophic nutrient C_*i*_ is given by:

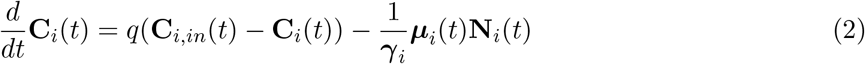

where *γ* is a vector of bacterial yield for each auxotrophic species with respect to their nutrient and **C**_*in*_ is a vector of the concentration of each nutrient flowing into the reactor (which is the quantity controlled by the reinforcement learning agent). Note that we assume all the auxotrophs are independent, i.e. each autotrophic nutrient is only used by one population.

The growth rates of each population is modelled using the Monod equation:

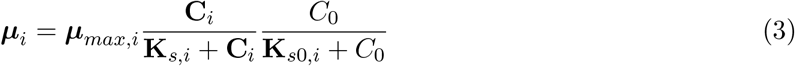

where ***μ***_*max*_ is a vector of the maximum growth rate for each species, **K**_*s*_ is a vector of half-maximal autotrophic nutrient concentrations and **K**_*s*0_ is a vector of half-maximal concentrations *C*_0_ for the shared carbon source. Finally, the growth rate for each population was determined as:

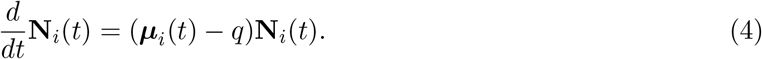

### 2.2 Reinforcement learning can be used to control the bioreactor system

Episodic Fitted Q-learning (Algorithm 2, Methods) was applied to a model of the system shown in Figure 1 (details in supplementary information) using the parameters in Table S1. This model was parametrised to simulate the growth of two distinct *E. coli* strains in a continuous bioreactor, with glucose as the shared carbon source, *C*_0_, and arginine and tryptophan as the auxotrophic nutrients *C*_1_ and *C*_2_. The reward was selected to penalize deviation from target populations of [*N*_1_, *N*_2_] = [20, 30] × 10^9^ cells L^−1^. Specifically, the reward function was: 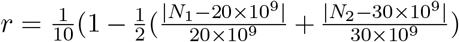. (The scaling was selected to ensure that the maximum possible reward was 0.1, which helped prevent network instability.) The agent was trained on thirty sequential episodes, this provided enough data for the agent to learn while not being prohibitive in terms of computational time. Each episode was twenty four hours long with sampling and subsequent action choice every five minutes. The explore rate was initially set to ∊ = 1 and decayed as ∊ = 1 − log_10_(*aE*) where *E* is the episode number, starting at 0, and *a* = 0.3 is a constant that dictates the rate of decay. A minimum explore rate of ∊ = 0 was set and was reached by the end of training. Figure 2A shows the training performance of twenty replicate agents, each trained over thirty episodes. The twenty agents converged to a mean final return of 27.4 with a standard deviation of 0.33. The theoretical maximum return is 28.8; all twenty agents were thus able to learn near optimal policies. The population curve in Figure 2B shows the system behaviour when under control of a representative agent trained in one of the replicates (for all twenty replicates see Figure S5). The population levels track the targets, with some jitter as expected with a bang-bang controller. Figure 2C shows the value function reached by this representative agent at the end of training, indicating its assessment of the total return from each point in state space. As expected, the value peaks at the target point. The corresponding state-action plot, Figure 2D, shows that the agent has adopted a simple, intuitive feedback law: add the specific nutrient needed by a strain when its population level is below the target and refrain from adding the nutrient if it is above the target. From these results, we conclude that reinforcement learning can be successfully applied to the chemostat system with a practical inter-sampling period of five minutes, as predicted in Figure S1 (details in the supplement).

**Figure 2:**
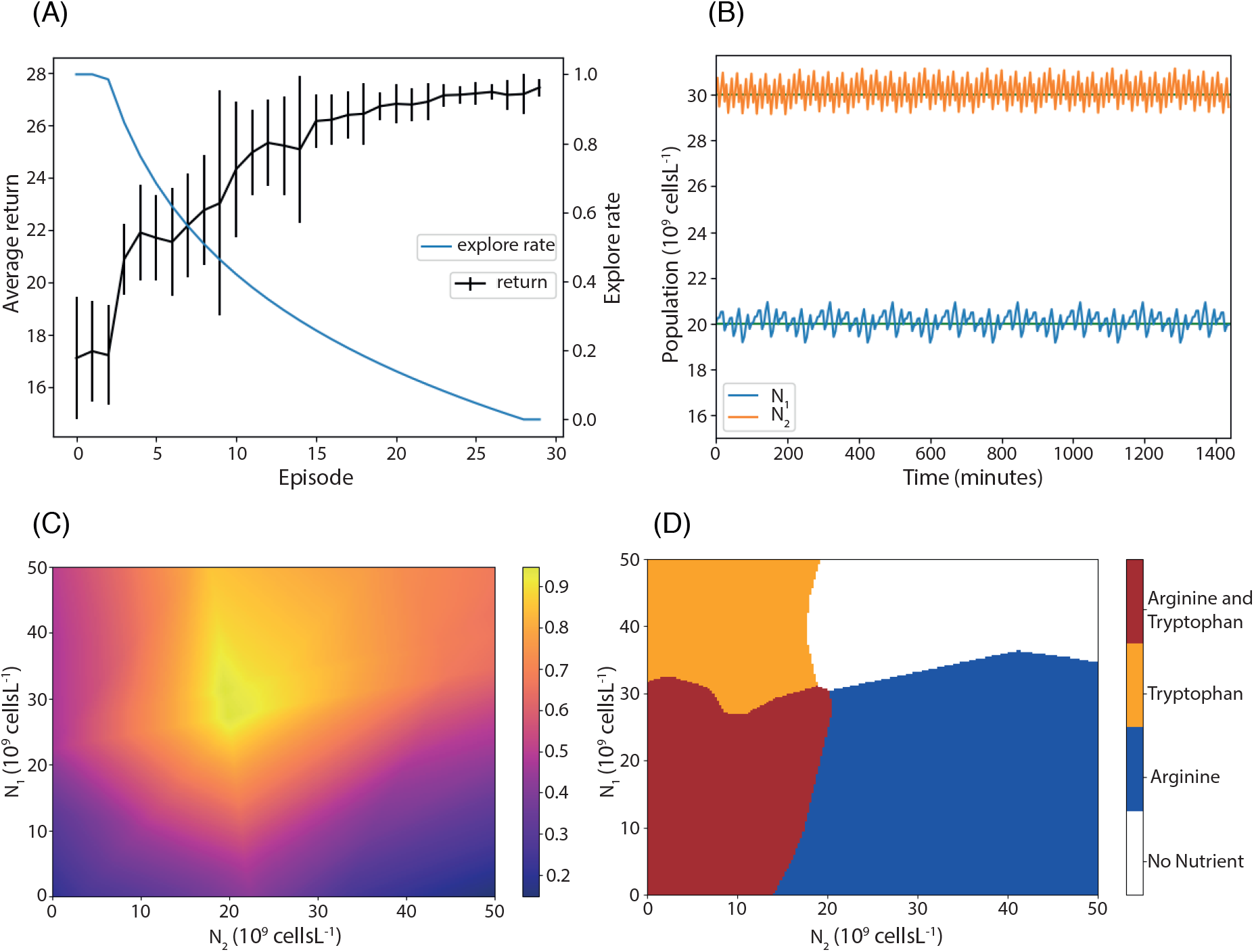
Reinforcement learning applied to the bioreactor system. (A) Performance of the agent improves and the explore rate decreases during training. The average of the return over twenty training replicates is plotted; error bars represent one standard deviation. (B) System behaviour under control of a trained agent for a twenty four hour period. Populations are maintained near target values (green lines). (C) Heatmap of the learned state value function; values are maximal at the target. (D) A learned state action plot.

### 2.3 Reinforcement learning outperforms proportional integral control for long sampling periods

As a comparison to a standard control approach, the reinforcement learning controller was compared to a traditional proportional integral controller. The controllers differ in that the proportional integral controller implements feedback over a continuous action space, whereas the reinforcement learning controller uses bang-bang control. For both controllers thirty episodes of data were generated, each twenty-four hours long, for a range of sampling-and-hold intervals: *t*_*s*_ = [5, 10, 20, 30, 40, 50, 60] mins by starting at initial state *S*_0_ = [20 × 10^9^ cells L^−1^, 30 × 10^9^ cells L^−1^, 0 *g L*^−1^, 0 *g L*^−1^, 1 *g L*^−1^] and sampling random input concentrations *C*_1_, *C*_2_ from [0, 0.1]*gL*^−1^. For each choice of sampling frequency, the reinforcement learning agent was trained using Fitted Q-iteration (Algorithm 1, Methods) on the dataset of thirty randomly generated episodes, while the proportional integral controller was tuned on an input-output model of the system derived from the same dataset (see Methods). The performance of the two controllers is illustrated in Figure 3, which shows how the performance depends on the choice of sampling frequency. For inter-sampling intervals longer than five minutes, the reinforcement learning controller outperforms the proportional integral controller. We conclude that reinforcement learning can produce comparable and even better performance, with the potential added advantage of a simpler implementation (the proportional integral controller employs continuous actions, whereas the reinforcement learning controller uses only bang-bang control). Moreover, for microbial chemostat systems that are difficult or expensive to sample at high frequency, reinforcement learning could be the preferred option.

**Figure 3:**
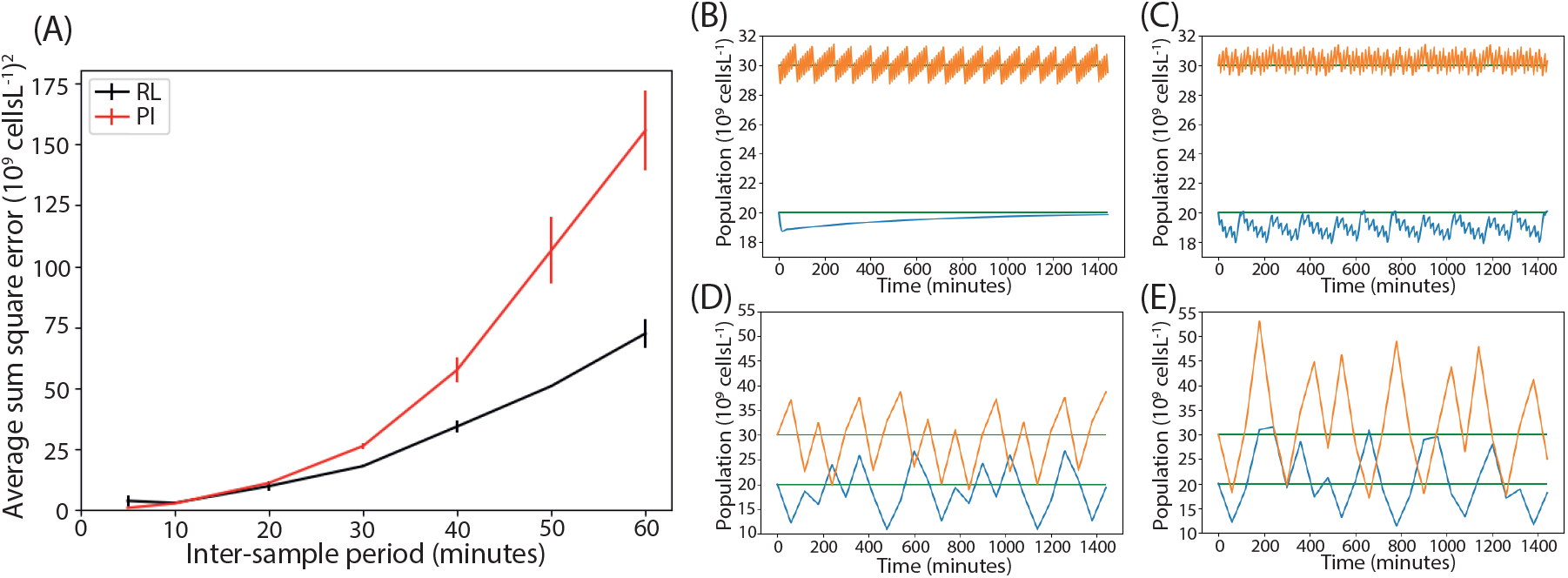
Comparison of reinforcement learning and proportional integral controllers. (A) The scaled average sum square error between the system state and the target. For long inter-sample periods, the reinforcement learning controller outperforms the proportional integral controller. The sum square error was calculated from population values that were scaled by a factor of 10^−9^. (B-C) Population time-courses under the proportional integral and reinforcement learning controllers respectively, with a five minute inter-sampling time. (D-E) Population time-courses under the proportional integral and reinforcement learning controller respectively, with a sixty minute inter-sampling time. Here the proportional integral controller allows the populations to stray further from their target values.

### 2.4 A good policy can be learned online using parallel bioreactors

A barrier to the use of reinforcement learning in real world applications is the amount of data required. We next show that Online Fitted Q-learning, a variant of Fitted Q-learning adapted to run in an online manner (Algorithm 3, Methods), can learn to control the chemostat system using an amount of data realistically obtainable in a single experiment. We trained an agent online on five chemostat models running in parallel. Each modelled the system described in Figure 1B and was run for the equivalent of twenty-four hours of real time. The agent took an action every five minutes, making an independent decision for each of the five chemostats from a single policy learned from experience gathered from all models. The reward was observed and the value function updated by the agent every ten time steps, using all experience gathered up to that time (Figure 4A). As in the previous sections, the initial microbial populations were set to the target value of [*N*_1_, *N*_2_] = [20, 30] × 10^9^ cells L^−1^. Figure 4B shows the online reward the agent received from the five chemostats. The initial reward was high, due to the initial populations being set to the target values. As the agent explored, the reward decreased and the standard deviation between the parallel chemostats increased because the agent took independent exploratory actions in each chemostat and drove them into different regions of state space. As time progressed, the reward from all five chemostats increased and the standard deviation decreased because the agent learned and moved all populations closer to the target. A pair of representative population time-courses is shown in Figure 4C (all five are shown in Figure S6). From these results, we conclude that Online Fitted Q-learning can be used to learn a policy in a data-efficient, online manner.

**Figure 4:**
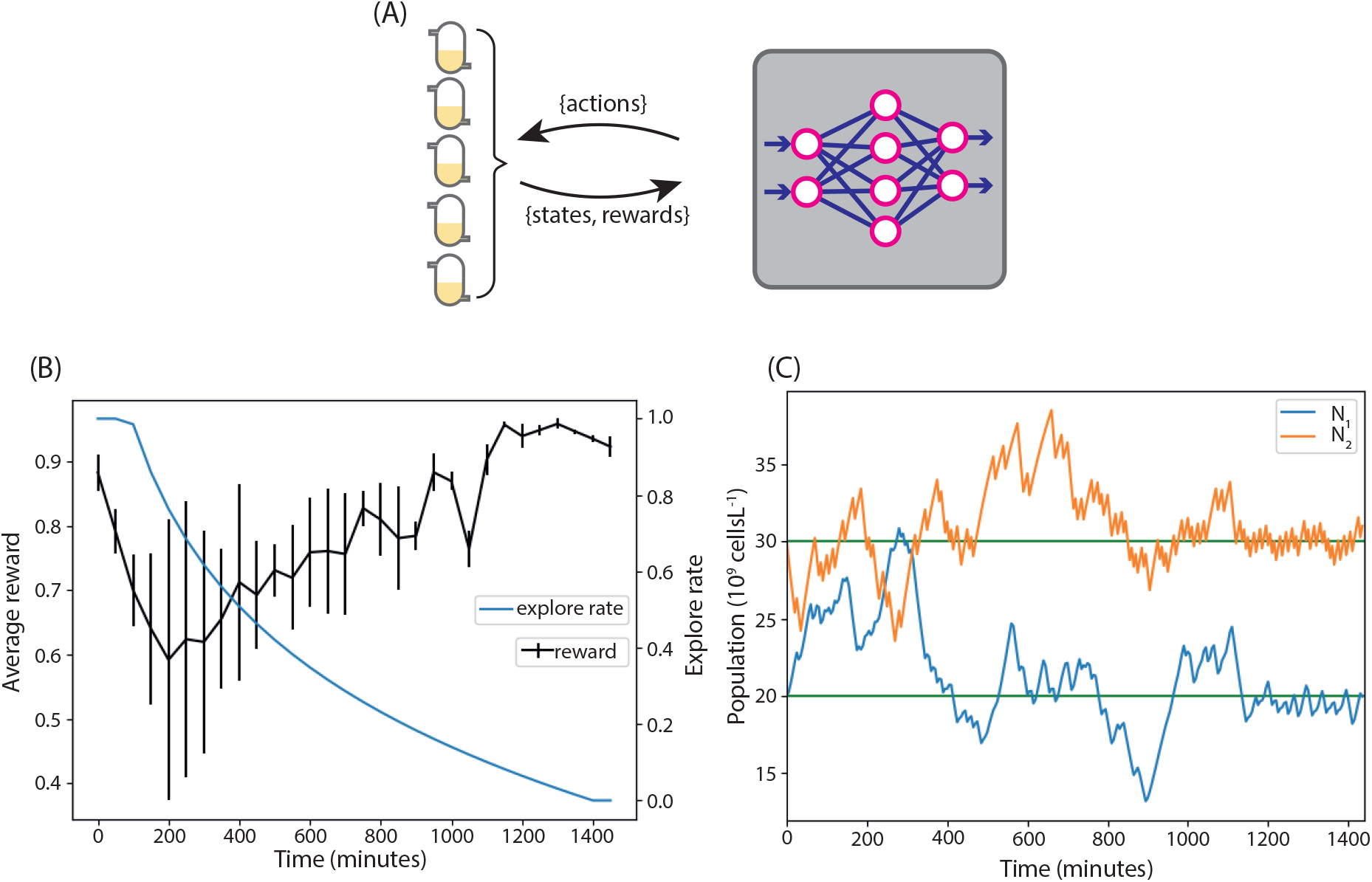
(A) A reinforcement learning agent was trained online on models of five parallel chemostats for twenty four hours. (B) The average reward received from the environments. By the end of the simulation all five chemostats were moved to the target population levels with very litte standard deviation in reward. (C) The population curve from one of the chemostats. During the exploration phase the population levels vary and random actions are taken, as the explore rate decreases they move to the target values.

### 2.5 The yield of a community-based product can be directly optimised

To demonstrate the ability of reinforcement learning to directly optimise the output of a communitybased bioprocess, the system in Figure 5A was modelled. Here, each microbial strain produces an intermediate product; *N*_1_ produces *A* and *N*_2_ produces *B*, each at a rate of 1 molecule per cell per hour. Factors *A* and *B* react to a product *P*, via the reaction 2*A* + *B* → *P*, which is presumed rapid. Consequently, the optimal state of the system has population ratio *N*_1_ : *N*_2_ = 2 : 1, with the populations at the maximum levels that the chemostat can support, which in our model means that all the carbon source, *C*_0_, is being consumed. In this case, we set the agent’s reward to be proportional to the amount of product produced by the bioreactor. We again take the observed state and the available actions to be the population levels and the bang-bang auxotrophic nutrient inflow rates, respectively. We set the initial populations to [*N*_1_, *N*_2_] = [20, 30] × 10^9^ cells L^−1^ as before. Ten replicate agents were trained using Episodic Fitted Q-learning (Algorithm 2, Methods). Performance in terms of the return is shown in Figure 5B. The average ratio of the population levels in steady state (the last 440 minutes of the simulation), over all agents, was 1.99 (with s.d. 0.08), showing convergence to near optimal populations in all replicates. A representative population time-course is shown in Figure 5C. Likewise, the average final concentration of the carbon source was 0.11% (s.d. 0.015%) of the source concentration, showing that in all cases the total population was close to the carrying capacity of the chemostat. As shown in Figure 5D, the replicates showed very little deviation in the final product output. However, in the initial phase of moving and stabilising the populations to the optimal levels, there is significant deviation. This suggests that most of the deviation in return shown in Figure 5B is due to this initial stabilising phase and not to the final phase the agents reached. From this analysis, we conclude that the reinforcement learning agent can learn to move the system to, – and keep it at – the near optimal state for product formation in a model free way.

**Figure 5:**
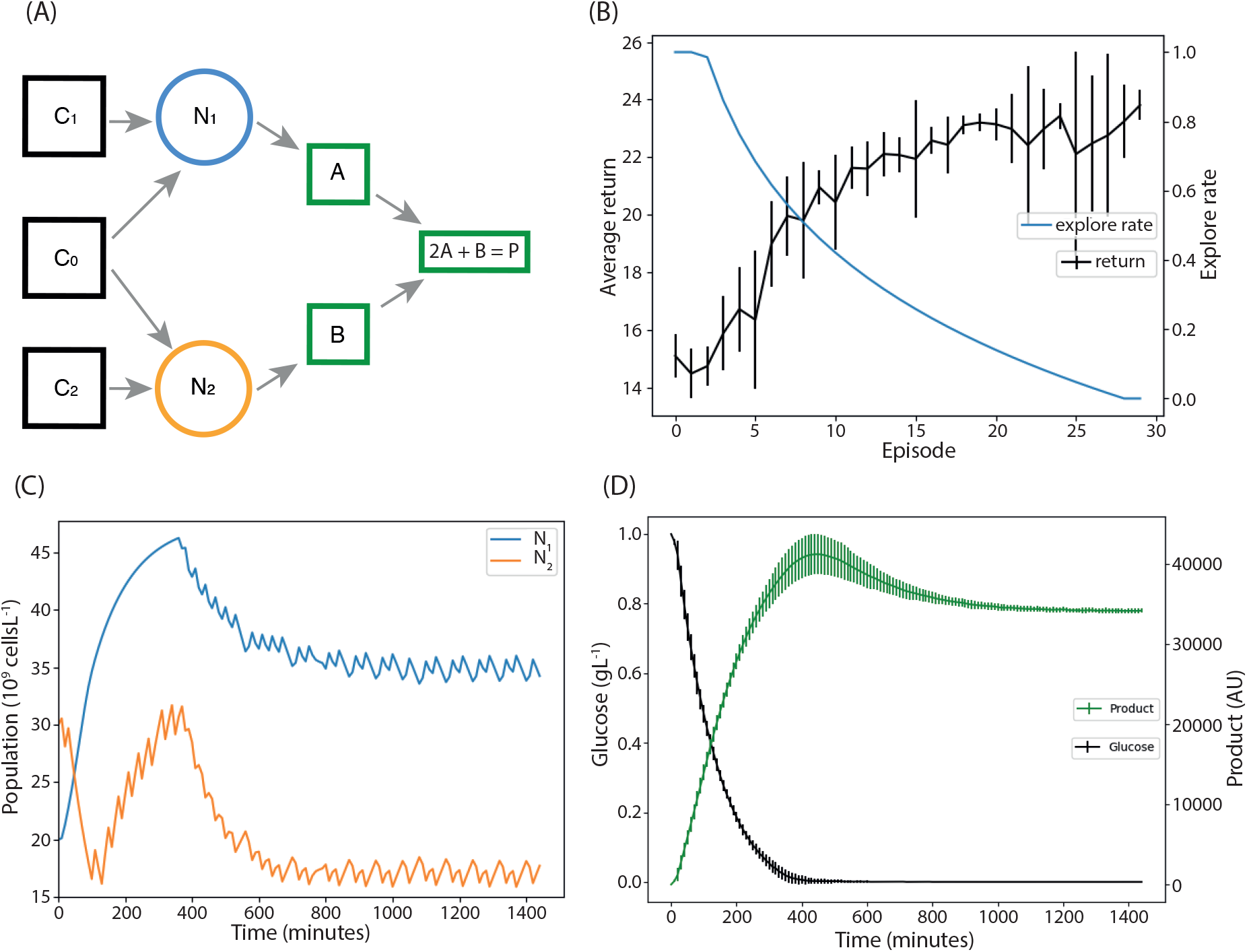
Using reinforcement learning to optimise product output. (A) Each microbial population produces an intermediate; these react to produce the desired product. (B) Training performance of ten reinforcement learning agents trained to optimise product output. (C) The resulting population curves of the system under control of a representative agent. The populations reach and then are maintained at the optimal level for product production. (D) The levels of carbon and product inside the chemostat. After the initial phase all carbon is being consumed. The levels of product peak as all initial carbon is used, then reach a level supported by the carbon supply to the reactor.

**Figure 6:**
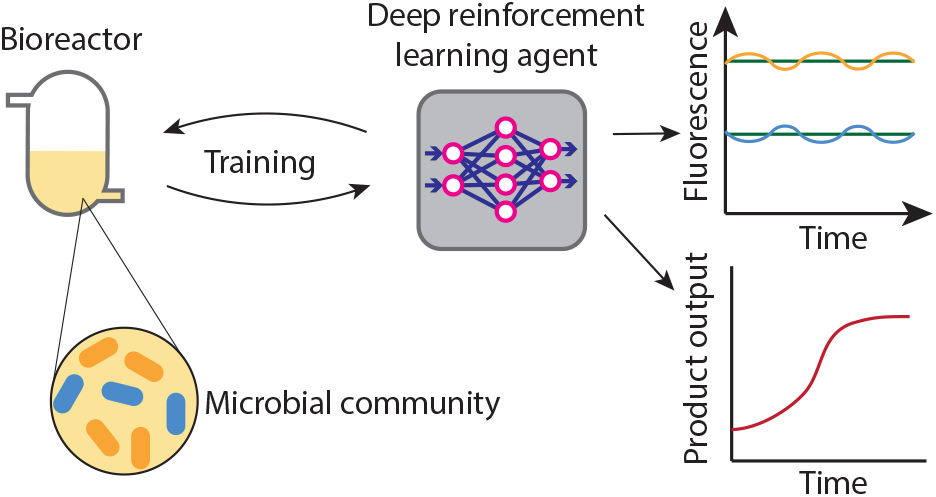
GRAPHICAL ABSTRACT

## 3 Discussion

We have applied deep reinforcement learning, specifically Neural Fitted Q-learning, to the control of a model of a microbial co-culture, thus demonstrating it’s efficacy as a model free control method that has the potential to compliment existing techniques. We have shown that reinforcement learning can perform better than the industry standard, PI control, when faced with long sample-and-hold intervals. In additional, it was shown that the data efficiency of Neural Fitted Q-learning can be used to learn a control policy in a practically feasible, twenty-four hour experiment. Reinforcement learning is most often used in environments where data is cheap and effectively infinitely available. Importantly, out work shows that it can also be realistically used to control microbial co-cultures. Finally it is shown that the output of a bacterial community can be optimised in a model free way using only knowledge of microbial population levels and the rate of product output, showing that industrial bioprocess optimisation is a natural application of this technique.

Overall, we have demonstrated the potential for control of multi-species communities using deep reinforcement learning. As synthetic biology and industrial biotechnology continue to adopt more complex processes for the generation of products from fine chemicals to biofuels, engineering of communities will become increasingly important. This work suggests that leveraging new developments in artificial intelligence may be highly suited to the control of these valuable and complex systems.

## 4 Methods

### 4.1 Neural Fitted Q-learning algorithm

A value function is learned which maps state action pairs to values. Here a state transition is defined as the tuple (*s*_*t*_, *a*_*t*_, *r*_*t*_, *s*_*t+1*_) specifying, respectively, the system state, action taken, and reward received at time *t*, and the state of the system at time *t* + 1. From a sequence of these state transitions a sequence of Q-learning targets is created according to:

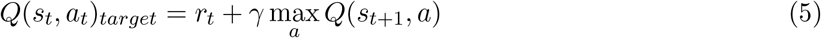

Here, the term max_*a*_ *Q*(*s*_*t+1*_, *a*), where *a* is an action that can be taken by the agent, gives an estimate of the total future reward obtained by entering state *s*_*t+1*_. This is weighted by *γ*, the discount factor, which dictates how heavily the possible future rewards weigh in on decisions. In this work, the discount factor was set to 0.9, which is a common first choice in reinforcement learning applications. A neural network is trained on the set of inputs {(*s*_*t*_, *a*_*t*_) ∀*t*} and targets {*Q*(*s*_*t*_, *a*_*t*_)_*target*_ ∀*t*} generated from all training data seen so far (Algorithm 1). In Episodic Fitted Q-learning this was done after each episode (Algorithm 2) while in Online Fitted Q-learning this was done after each update interval (Algorithm 3). The number of Fitted Q-iterations was set to 10 for all Episodic Fitted Q-learning and was chosen depending on the amount of data in the agents memory for Online Fitted Q-learning. See supplement for further details (Figures S1-S3).

**Algorithm 1.**
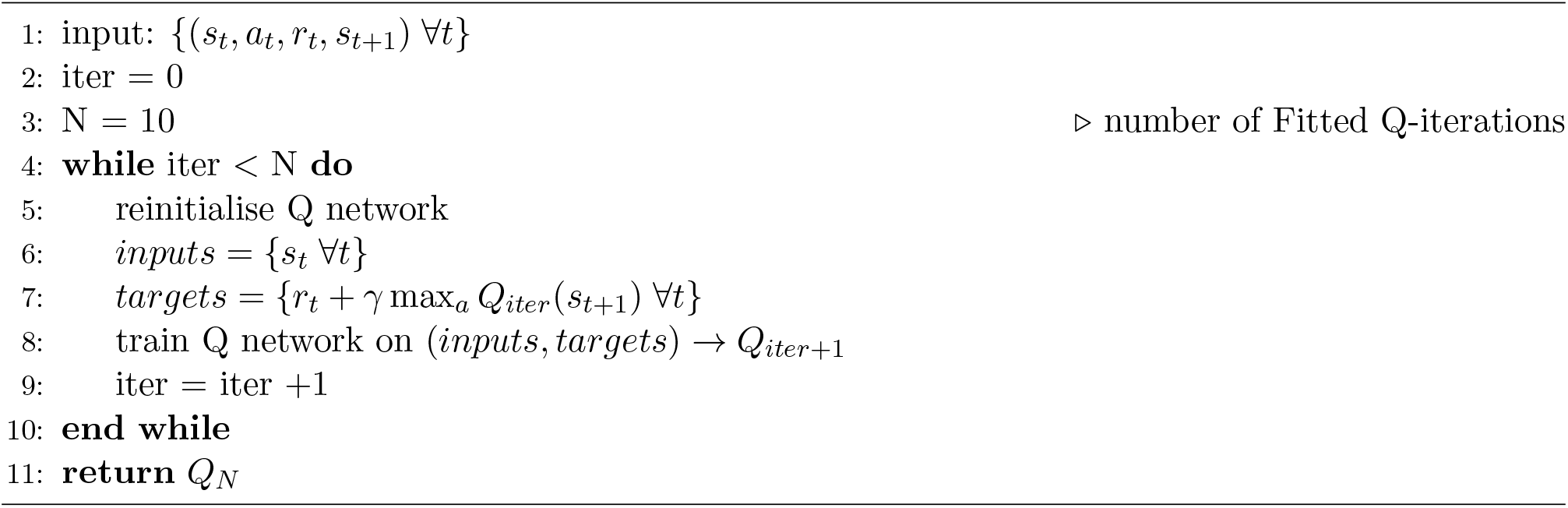
Fitted Q-iteration

We use an ∊-greedy policy in which a random action is chosen with probability ∊ and the action *a* = max_*a*_ Q(*s_t_, a*) is chosen with probability 1 − ∊. The *explore rate* ∊ was set to decay exponentially as training progressed. ∊-greedy is a widely used policy that has been proven effective [34, 29] and is easy to implement.

The state variables considered by the algorithm were the continuous populations of each species of microbe. The agent acted as a bang-bang controller with respect to each input nutrient, giving 2^*n*^ possible actions, where *n* is the number of nutrients. (In tis work, *n* = 2.)

The neural network that was used to estimate the value function consisted of two hidden layers of 20 nodes, following the approach in previous work [27]. Each node in the hidden layers used the ReLU activation function. The input layer had *n* nodes, one for each microbial strain; the linear output layer had 2^*n*^ nodes, one for each available action. We used the The Adam optimiser [35], because of its ability to dynamically adapt the learning rate, which is favourable when implementing reinforcement learning with a neural network [36]. The populations levels were scaled by a factor of 10^−5^ before being entered into the neural network; this generated values between 0 and 1 and prevented network instability.

**Algorithm 2.**
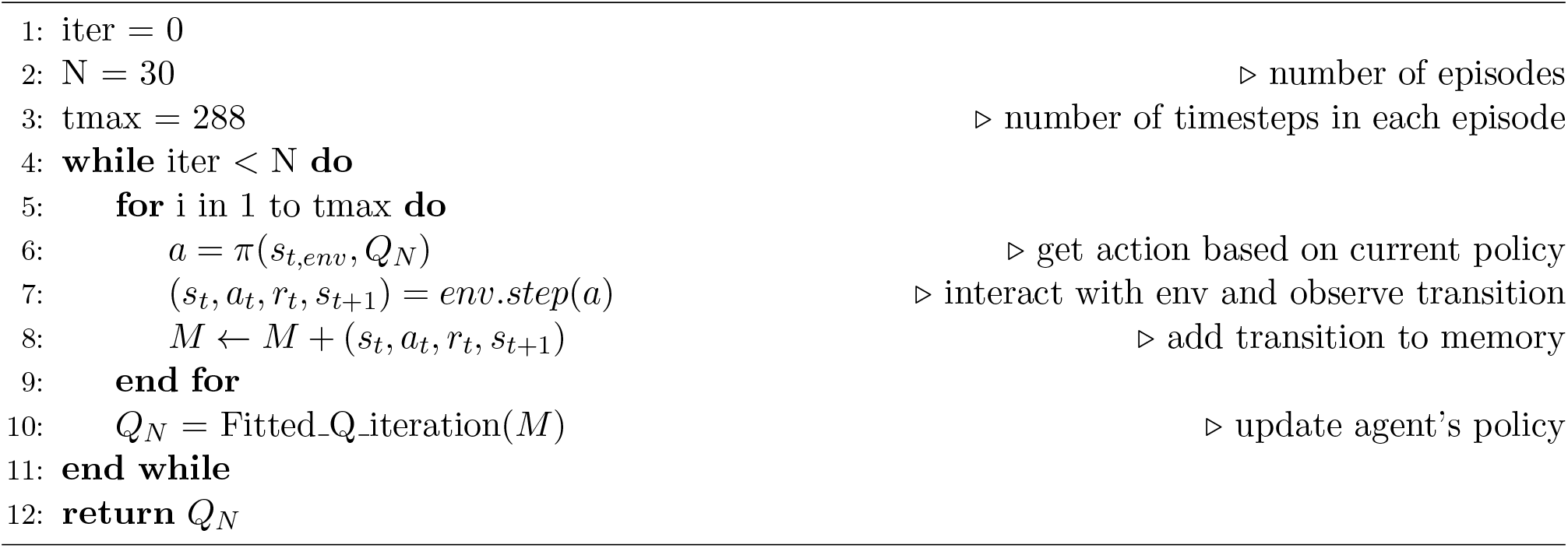
Episodic Fitted Q-learning

**Algorithm 3.**
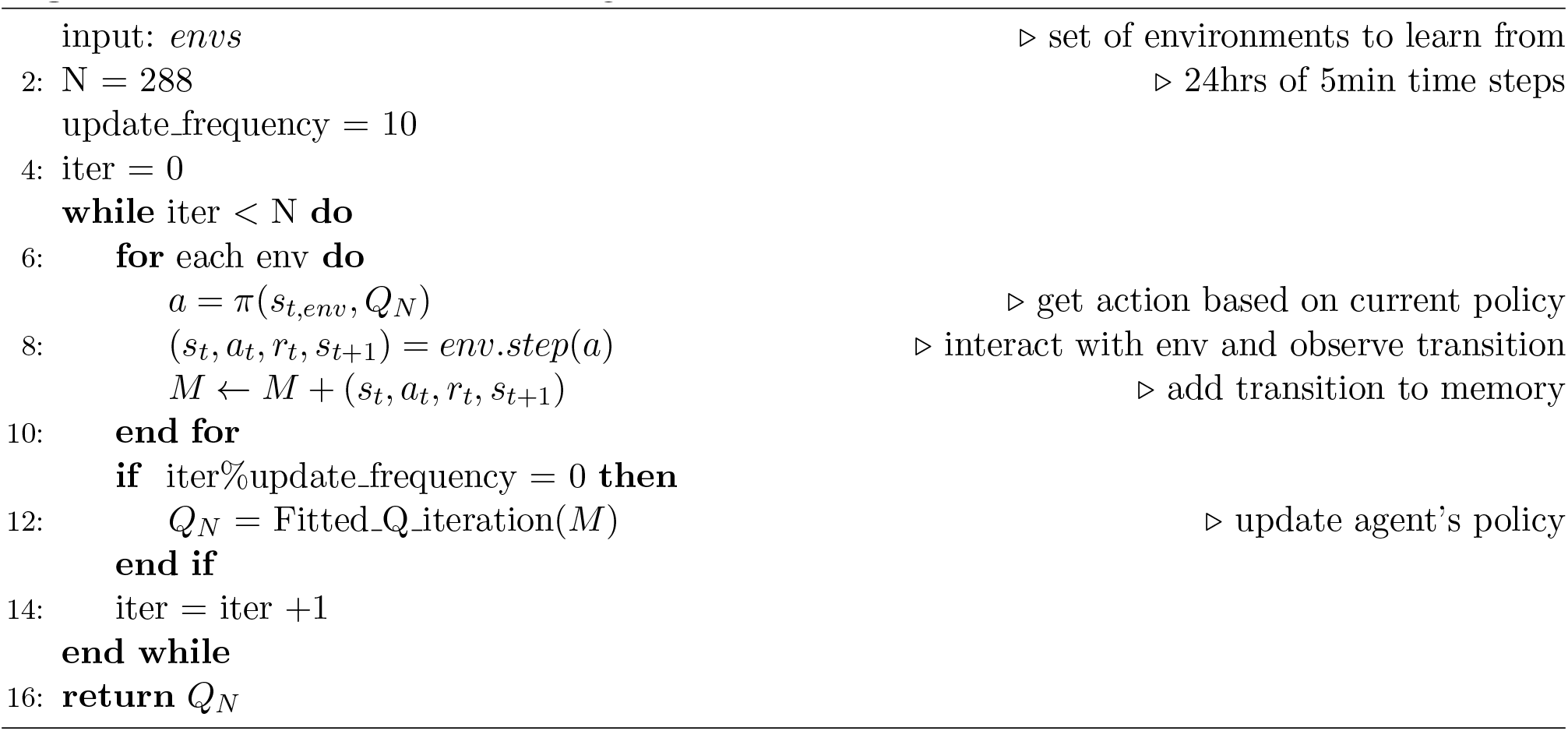
Online Fitted Q-learning

Python version 3.6.7 was used for all reinforcement learning code, available at http://www.python.org. The odeint function of SciPy (version 1.3.1) [37] was used to numerically solve all differential equations. The neural network was implemented in Google’s TensorFlow (version 1.13.1) [38]. Numpy (version 1.16.14) was used throughout [39]. The code and examples are available on GitHub: https://github.com/ucl-cssb/ROCC

### 4.2 Proportional integral controller tuning

For the comparison between reinforcement learning and PI control, we tested a range of sample-and-hold intervals ([5, 10, 20, 30, 40, 50, 60] mins). For each choice of sampling interval, we generated thirty, twenty-four hour long episodes, each starting from initial state *S*_0_ = [20000 × 10^6^ cells L^−1^, 30000 × 10^6^ cells L^−1^, 0 *g L*^−1^, 0 *g L*^−1^, 1 *g L*^−1^] and selecting actions randomly from [0, 0.1] *g L*^−1^. These thirty episodes were used as training data for the Fitted Q-agent. From each dataset, an input-output model was constructed using the plant identification function in the PID tuner app of MATLAB’s Simulink toolbox, which allows the identification of an input-output model for any input-output dataset. Here, the randomly chosen actions were used as input and the resulting populations (scaled by a factor of 10^−10^) were taken as output. The model was a state space model, of order chosen by the system identification app to best fit the data. The Akaike’s Final Prediction Error (FPE) of the model fits was of the order 10^−2^ for the 5 min sample-and-hold intervals, rising to a maximum of almost 1 for 60 min intervals (see supplementary file PI data.ods for full results). An independent input-output model was derived for each microbial population. These were used to tune two independent PI controllers, one controlling each population. We used independent controllers because the PI tuner app is only compatible with single input, single output systems. We considered a range of tuning objectives to assess the merits of tuning to minimise settling time, rise time or overshoot percentage. We found that minimising rise time led to high overshoot errors, while minimising overshoot percentage also led to high errors because the controller would be slow to reach the target. Tuning the controller to minimise settling time worked best for all cases tested and can be seen as a compromise between speed of response and robustness. Hence, for all results presented, the PI controllers were tuned to minimise settling time. All results for the PID tuning, including the gains, FPE, settling times, rise times and overshoot percentages can be found in the supplementary file PI data.ods. The Simulink diagram of the system is shown in Figure S8.

