## Supplementary Information for "Deep reinforcement learning for the control of microbial co-cultures in bioreactors"

October 29, 2019

### 1 Reinforcement learning parameter tuning

We carried out preliminary investigations to calibrate parameters for the reinforcement learning controller, as follows.

#### 1.1 Minimum inter-sampling period

The theoretical convergence guarantees of reinforcement learning assume that it is applied to a Markov decision process [1]. The two-strain chemostat system we use here has five state variables: two autotrophic nutrient concentrations, the concentration of carbon source and the two microbial population levels. Only the population levels are known to the agent, meaning the system is only partially observed and is hence not a Markov decision process. There are methods to extend reinforcement learning to partially observed Markov decision processes, including incorporating time series information using a recurrent network [2], keeping track of approximate belief states of the hidden variables [3] or using Monte-Carlo methods [4]. To assess whether these computationally expensive methods would be required, we determined, as follows, the minimum sample-and-hold interval that allowed the agent to accurately predict the reward resulting from a chosen action. (Intuitively, this can be thought of as the minimum sample-and-hold interval in which an action has time to have an effect on the observed states (the population levels).) To determine this minimum interval length, we first generated system trajectories of $(s_t, a_t, r_t)$ resulting from random actions. The agent was trained on these sequences to predict the reward $r_t$ from the state action pairs $(s_t, a_t)$. We repeated this process one hundred times for each of the following sampling times: [1,2,3,4,5,10] minutes. The results, shown in Figure S1A, indicate that at time steps lower than four minutes the agents are unable to accurately predict the reward received from the state and action, meaning reinforcement learning cannot be effective. However at four minutes and above the reward prediction is accurate. We concluded that by using intervals of four minutes or longer, the sophisticated non-Markovian methods mentioned above would not be required for this application. For most of the results in the main text a time step of five minutes was used. Figure S1B shows the reward prediction for both one- and five-minute time steps, showing that the agent performs well for five minutes intervals and poorly for one minute intervals.

*
†

##### 1.2 Number of Fitted Q-iterations to avoid over fitting

To determine how many Fitted Q-iterations to implement, we generated sequences of $(s_t, a_t, r_t)$ of varying lengths and trained Fitted Q-agents to predict the rewards. We determined the training and testing error for each Fitted Q-iteration, with a maximum of 40 iterations. Figure S2 shows the results of repeating this process 100 times for each sequence length. The data reveal clear over fitting for the datasets shorter than 200 time steps long and a reduction in testing error as the sequence length increases (i.e. with more training data). For each sequence length, the training process with 4 fitted Q iterations gave the smallest testing error (except for 100 training timesteps, where 5 iterations performed marginally better). With a training set of 200 time steps, no significant overfitting occurred.

##### 1.3 Number of Fitted Q-iterations for value convergence

Another consideration is how many Fitted Q-iterations are required for the values to converge via bootstrapping. For this analysis, we generated 100 sequences of $(s_t, a_t, r_t)$, each one thousand time steps long. For each sequence, the actual values were calculated and Fitted Q-iteration was used to obtain predicted values. After each Fitted Q-iteration, the error between the predicted and actual values was recorded. As shown in Figure S3, the values convergence after about ten iterations. Using this and the information from the previous section the number of Fitted Q-iterations was chosen depending on the length of the agents memory to both prevent overfitting and to allow convergence via bootstrapping. For all episodic Q-Learning the number of Fitted Q-iterations was set to 10 as the agents memory always contains at least one episode of 288 transitions. For online Q-Learning the number of Fitted Q-iterations was set to 4 if there were less than 100 transitions in the agents memory, 5 if there were 100-199 transitions and 10 if there were 200 or more transitions.

#### 2 Parameter table

| Parameter | Description | Value | Unit | Source |
|---|---|---|---|---|
| $\mathbf{C}_{0,in}$ | Reservoir concentration of carbon source | 1 | g L$^{-1}$ | Experimentally controllable |
| q | Flow rate | 0.5 | h$^{-1}$ | Experimentally controllable |
| $\boldsymbol{\gamma}_0$ | Yield coefficient for common substrate | $4.8 \times 10^{11}$ | cells g$^{-1}$ | [5] |
| $\boldsymbol{\gamma}_1$ | Yield coefficient for arginine | $5.2 \times 10^{11}$ | cells g$^{-1}$ | [6] |
| $\boldsymbol{\gamma}_2$ | Yield coefficient for tryptophan | $4.4 \times 10^{11}$ | cells g$^{-1}$ | [6] |
| $\boldsymbol{\mu}_{max,1}$ | Maximum growth rate | 1 | h$^{-1}$ | [7] |
| $\boldsymbol{\mu}_{max,1}$ | Maximum growth rate | 1.1 | h$^{-1}$ | [7] |
| $\mathbf{K}_{s,0}$ | Saturation constant for the carbon source | $6.845928 \times 10^{-5}$ | g L$^{-1}$ | [6] |
| $\mathbf{K}_{s,1}$ | Saturation constant for arginine | $4.9 \times 10^{-4}$ | g L$^{-1}$ | [6] |
| $\mathbf{K}_{s,2}$ | Saturation constant for tryptophan | $1.02 \times 10^{-7}$ | g L$^{-1}$ | [6] |

Table 1: Double auxotroph system. Parameter values used for simulations of a system consisting of two auxotrophic populations of bacteria with competition for nutrients. $\boldsymbol{\mu_{max}}$ values were chosen using values from the literature [7] as a guide.

#### 3 Supplementary figures

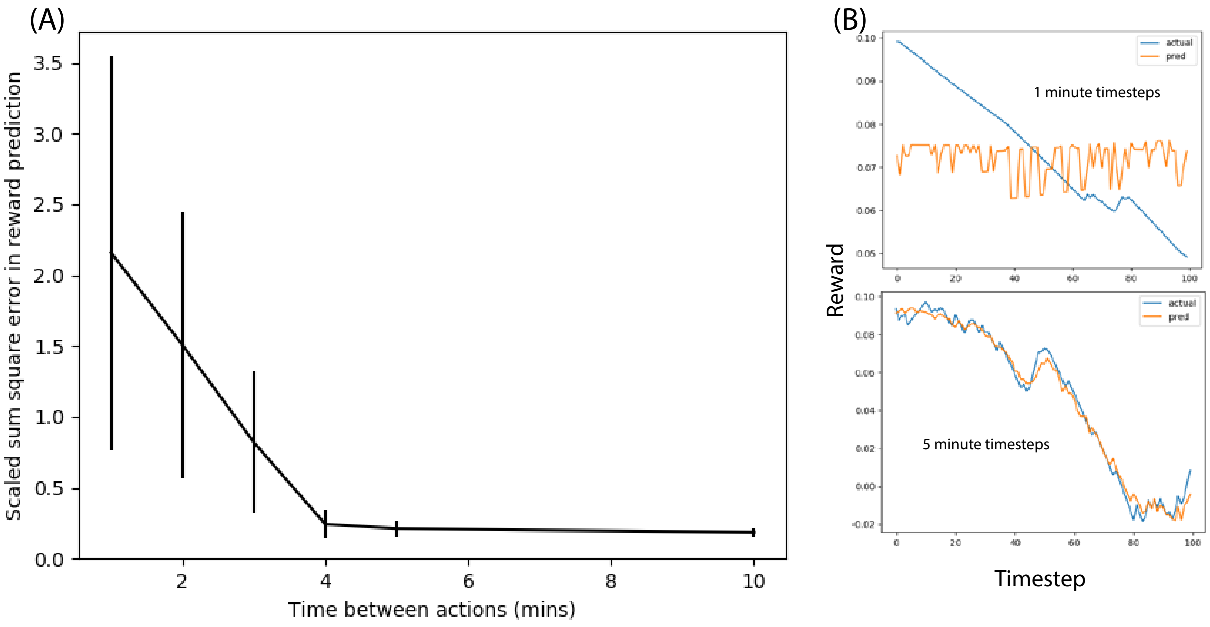


Figure 1: Identifying the minimum timestep above which the chemostat system behaves as a Markov decision process. (A) The error in reward prediction is negligible for time steps above four minutes. Error bars represent one standard deviation. (B) The predicted vs actual reward for one minute and five minute timesteps. Markov decision-based learning is not possible for the short one-minute intervals, but performs well for five-minute intervals.

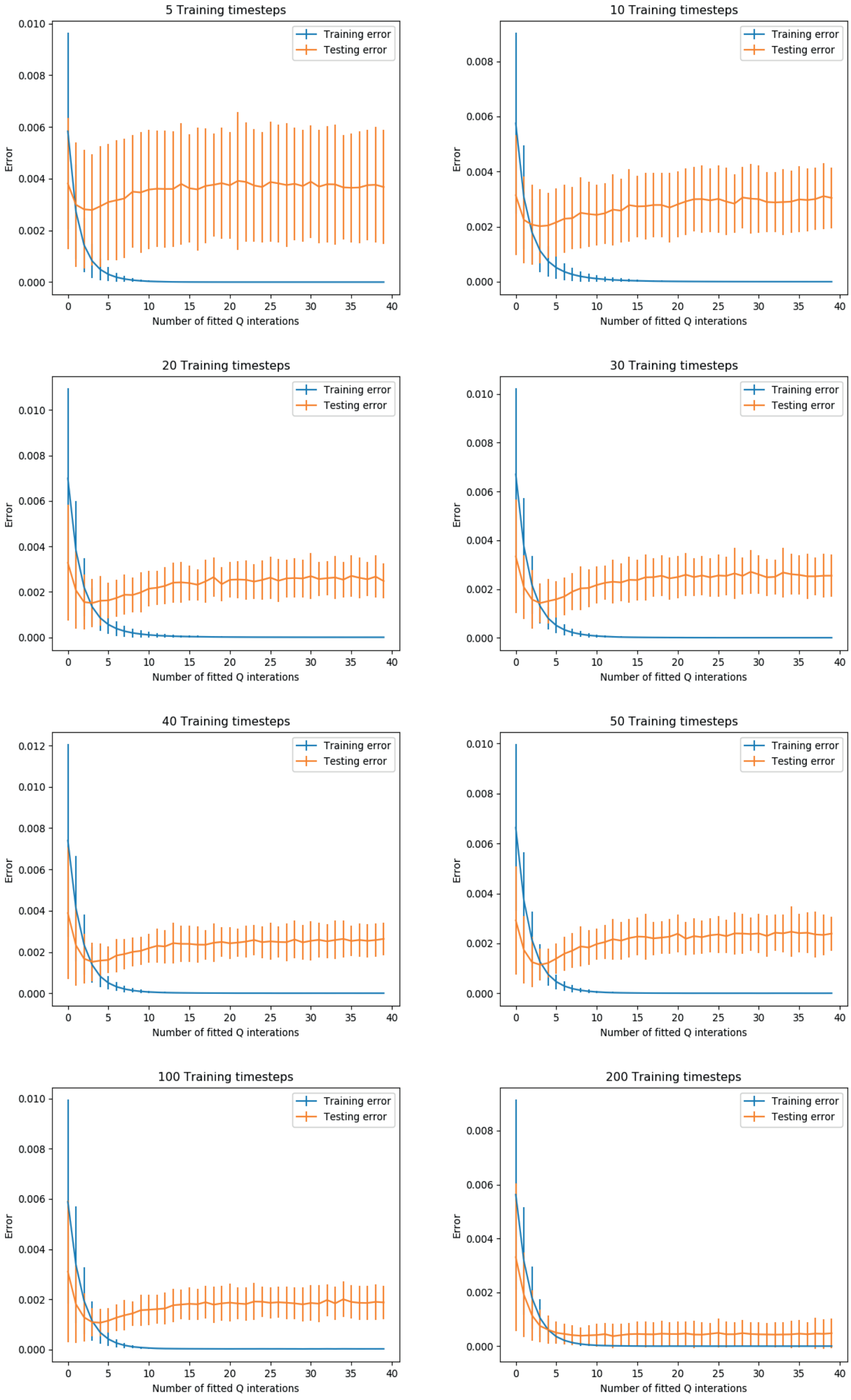


Figure 2: The training (blue) and testing (orange) accuracy of a Fitted Q-agent to predict rewards from states and actions was tested after every Fitted Q-iteration. Overfitting is seen for number of transitions less than 200. Error bars represent one standard deviation.

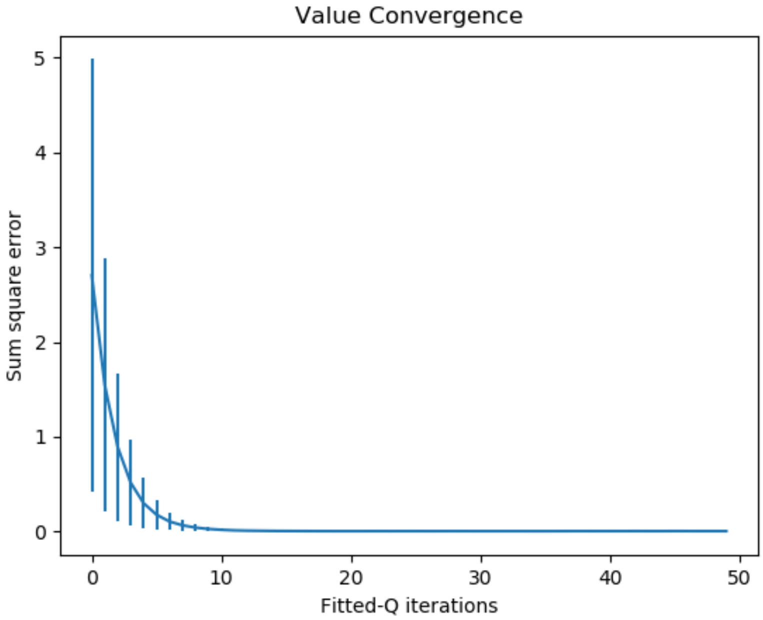


Figure 3: The scaled error between actual and predicted values as Fitted Q-iterations are done

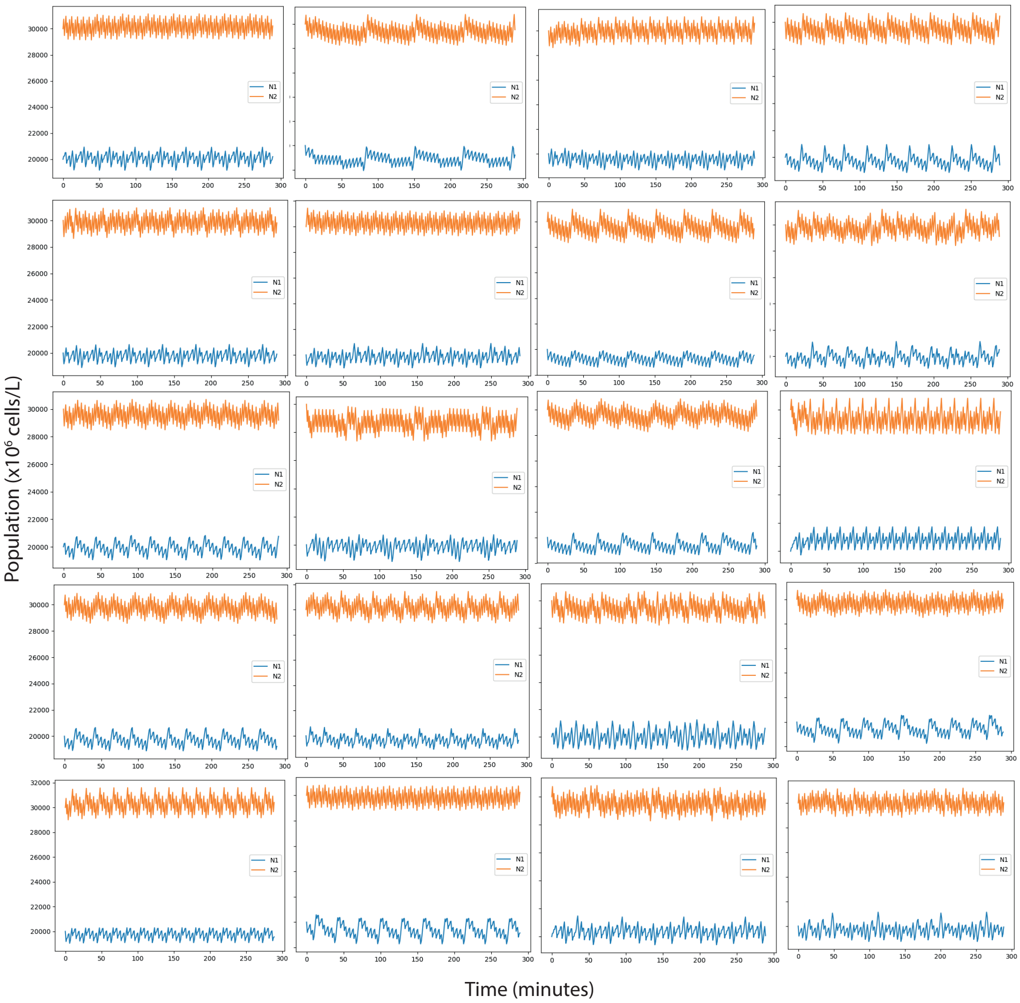


Figure 4: Population curves of twenty trained agents controlling the chemostat system

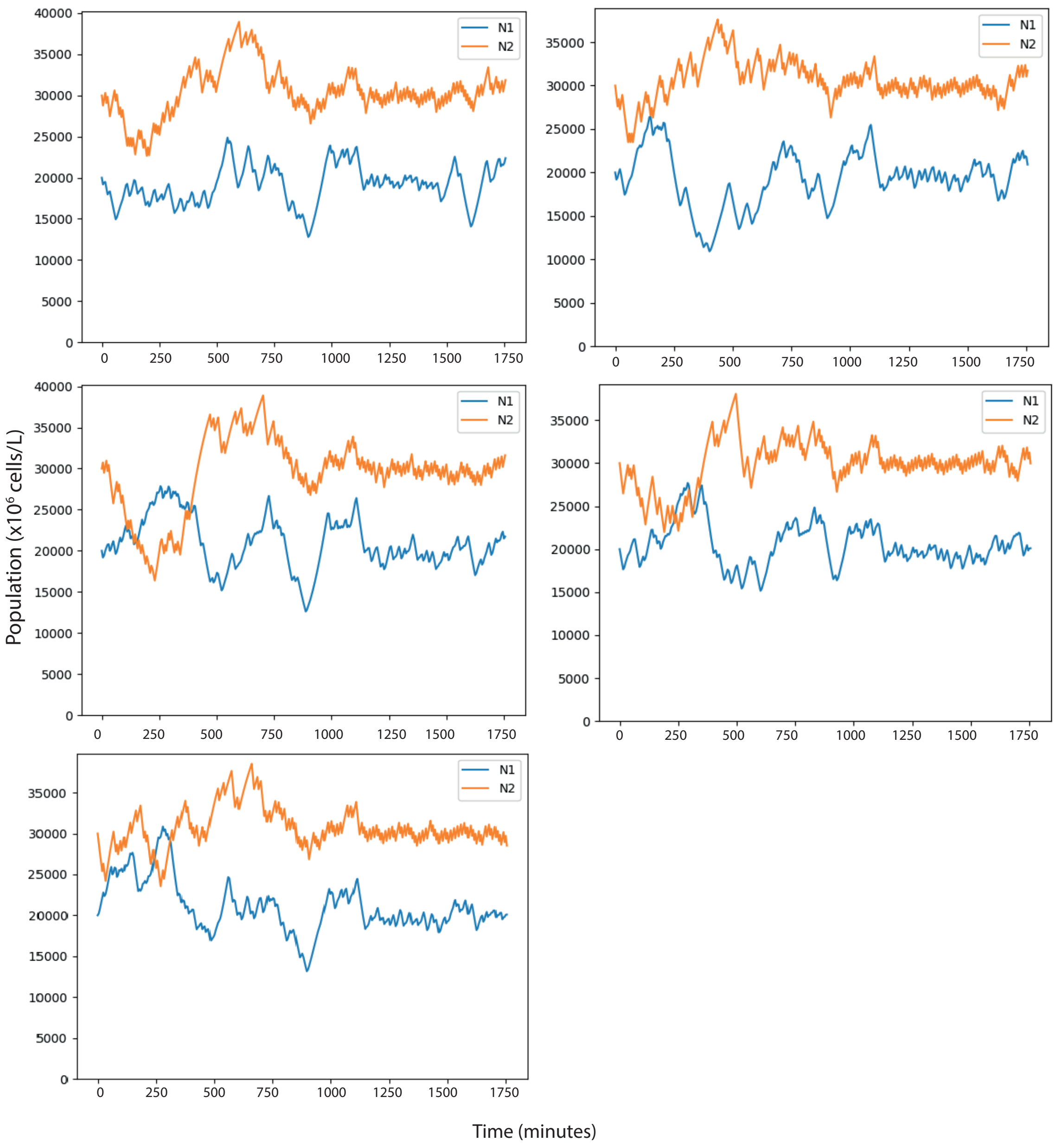


Figure 5: Populations curves of five chemostats running in parallel while under online control of a single agent. Here the agent is trained for 1440 minutes (twenty-four hours) and then allowed to control the system for a further 310 minutes to show that the target system behaviour is maintained.

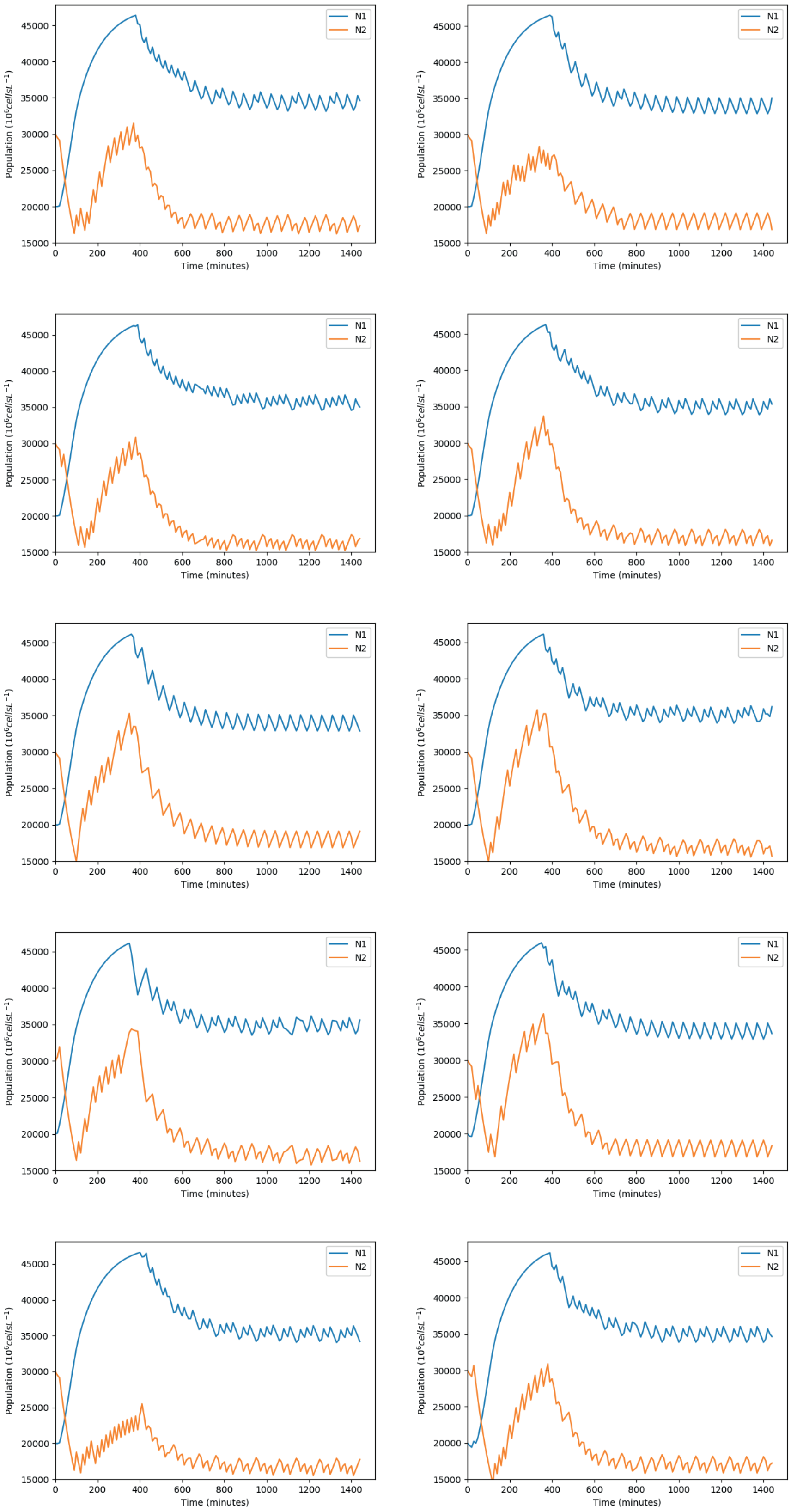


Figure 6: Population curves of ten trained agents controlling the chemostat system with the goal of optimising product output

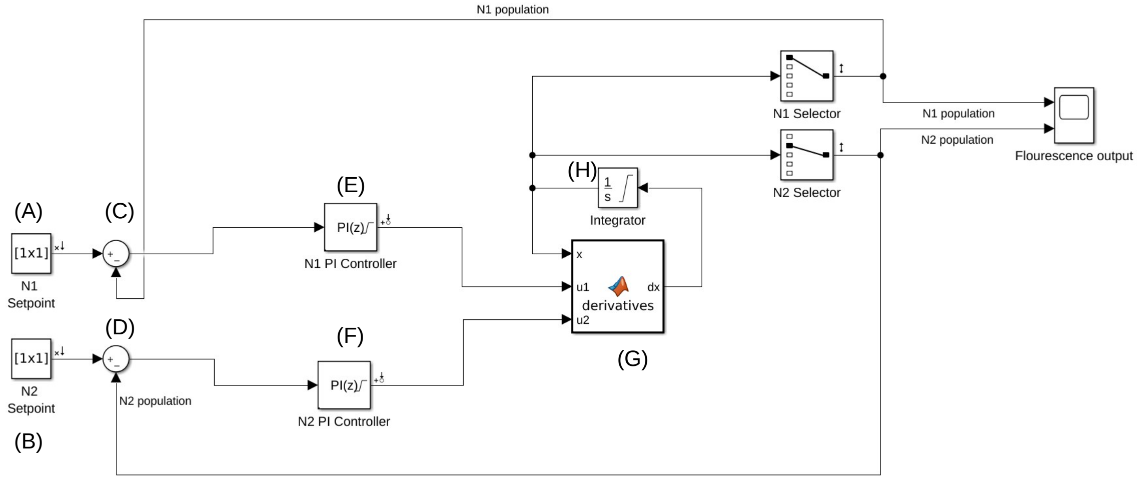


Figure 7: The Simulink diagram of the system and PI controllers. (A,B) the setpoints or target population levels, from which the error is calculated (C,D) and used by the PI controllers (E,F) to adjust nutrient levels. The system of ODEs (G) is solved by a continuous time integrator (H).
